# GRIDSS2: comprehensive characterisation of somatic structural variation using single breakend variants and structural variant phasing

**DOI:** 10.1101/2020.07.09.196527

**Authors:** Daniel L. Cameron, Jonathan Baber, Charles Shale, Jose Espejo Valle-Inclan, Nicolle Besselink, Arne van Hoeck, Roel Janssen, Edwin Cuppen, Peter Priestley, Anthony T. Papenfuss

**Affiliations:** Bioinformatics Division, Walter and Eliza Hall Institute of Medical Research, Parkville, Australia; Department of Medical Biology, University of Melbourne, Australia; Hartwig Medical Foundation Australia, Sydney, Australia; Hartwig Medical Foundation, Science Park 408, Amsterdam, The Netherlands; Center for Molecular Medicine and Oncode Institute, University Medical Center Utrecht, Heidelberglaan 100, Utrecht, The Netherlands; Peter MacCallum Cancer Centre, Melbourne, Australia; Sir Peter MacCallum Department of Oncology, University of Melbourne, Australia

**Keywords:** Single breakends, somatic, structural variation

## Abstract

GRIDSS2 is the first structural variant caller to explicitly report single breakends - breakpoints in which only one side can be unambiguously determined. By treating single breakends as a fundamental genomic rearrangement signal on par with breakpoints, GRIDSS2 can explain 47% of somatic centromeric copy number changes using single breakends to non-centromeric sequence, with chromosome 1 exhibiting a unique centromeric rearrangement signature. On a cohort of 3,782 deeply sequenced metastatic cancers, GRIDSS2 achieved an unprecedented 3.1% false negative rate and identified a novel 32-100bp duplication signature. Somatic structural variants are highly clustered with GRIDSS2 phasing 16% using just paired-end sequencing.

## Background

The reliable detection of structural variants (SVs) is critical to understanding the role genome architecture plays in health and disease. This is especially important in cancer and precision medicine where structural variation can be a key driver mutation ^1,2^. Over the past decade, many tools have been developed for the detection of genomic rearrangements, which have been the subject of recent extensive benchmarks ^3,4^. These tools fall broadly into two camps: those that detect changes in DNA abundance, known as copy number variant or aberration (CNV/CNA) callers, and those that detect non-reference DNA adjacencies, known as structural variant (SV) or breakpoint callers. While CNAs and SVs are merely two different viewpoints of the underlying genomic rearrangements, the methods of detection are fundamentally different. Here, we address the problem of SV detection and show that breakpoint detection alone is insufficient for the comprehensive characterisation of somatic genomic rearrangements that occur in cancer. A third genomic rearrangement primitive is essential: single breakends.

The Variant Call Format (VCF)^5^ defines a single breakend as a breakpoint in which only one side can be unambiguously placed. This can occur due to one of two reasons. Firstly, the sequence on one side of the breakpoint could be absent from the reference. Either non-reference sequence could be present due to the integration of foreign DNA (e.g. provirus) or the reference could lack sequence present in the sample. Secondly, breakpoints into highly repetitive regions cannot be unambiguously placed. Single breakends allow the representation of such breakpoints. Such rearrangements are common in cancer and by reporting single breakends the rearrangement landscape of regions previously considered inaccessible to short read sequence can be explored.

Short read-based SV detection algorithms identify breakpoints by finding clusters of reads that do not support the reference allele. Typically these use discordant read pairs ^6^, or split reads^7^, with some callers also considering reads with unmapped mates ^8^ and soft-clipped reads ^9^. More sophisticated callers incorporate assembly either through de novo assembly ^10^, targeted breakpoint assembly ^11^, or breakend assembly ^12^. These callers report breakpoints, that is, novel adjacencies. When reads cannot be unambiguously mapped on either side, a breakpoint call cannot be made and information is lost. Some callers have attempted to address this by considering multiple alignment locations for each read ^13^ but this only works for regions with a small number of potential alignment locations and has proven impractical for general use. Single breakend calling has the potential to improve short read caller sensitivity above the 50% reported in recent benchmarking^3–5^.

As we move closer to a world in which the CNA and SV primitives can be reliably detected, accurate interpretation of the causative biological events becomes increasingly possible by integrated analysis of this knowledge. While progress has been made on derivative chromosome reconstruction using long reads ^14^, reconstruction of complex events such as chromothripsis has been problematic for short reads ^15,16^. To date, SV phasing has been used to reduce the complexity of reconstruction for long read based approaches ^17^ but has not been done by short read callers. The ability of phase somatic structural variants is limited by the read length and, for short read data, by the library fragment size - typically less than 500bp.

Here, we demonstrate the power of single breakend variant calling using GRIDSS2 - a somatic structural variant caller that reports single breakends and phases nearby structural variants. Running GRIDSS2 on 3,782 metastatic solid tumours with matched normal samples from the Hartwig cohort we show that, due to the high prevalence of somatic breakpoints involving low-mappability sequences, GRIDSS2 achieves a false negative rate lower than possible with a traditional breakpoint-only caller. The precision and sensitivity of GRIDSS2 in conjunction with single breakend variant calling and SV phasing lay a strong foundation for downstream tools that enable a deeper understanding of the nature of somatic genomic rearrangements.

## Results

GRIDSS2 utilises the same high-level approach as the first version of GRIDSS, assembling all reads that potentially support a structural variant using a positional de Bruijn graph breakend assembly algorithm^12^. Breakend contigs are then realigned back to the reference to identify breakpoints and probabilistic structural variant calling is performed based on both the aligned reads and assembled contigs. Single breakend variant calling uses the same probabilistic variant calling approach as breakpoint calling, but instead of split reads, discordant read pairs, and assembly contigs with chimeric alignments support, single breakends are called based on soft-clipped reads, reads with unmapped or ambiguously mapping mates, and assemblies with unmapped or ambiguously mapping breakend sequence (Figure 1a). SV phasing is performed based on assembly contigs and the presence of transitive calls (Figure 1b). SVs are phased cis if an assembly spans both breaks or a transitive call is found, and phased trans if an assembly involves one SV but supports the reference at the other. Since assembly contig length is limited by the library fragment size only nearby SVs can be phased. GRIDSS2 includes a 16-step somatic filter specifically tuned for deeply sequenced tumour/normal samples.

**Figure 1.**
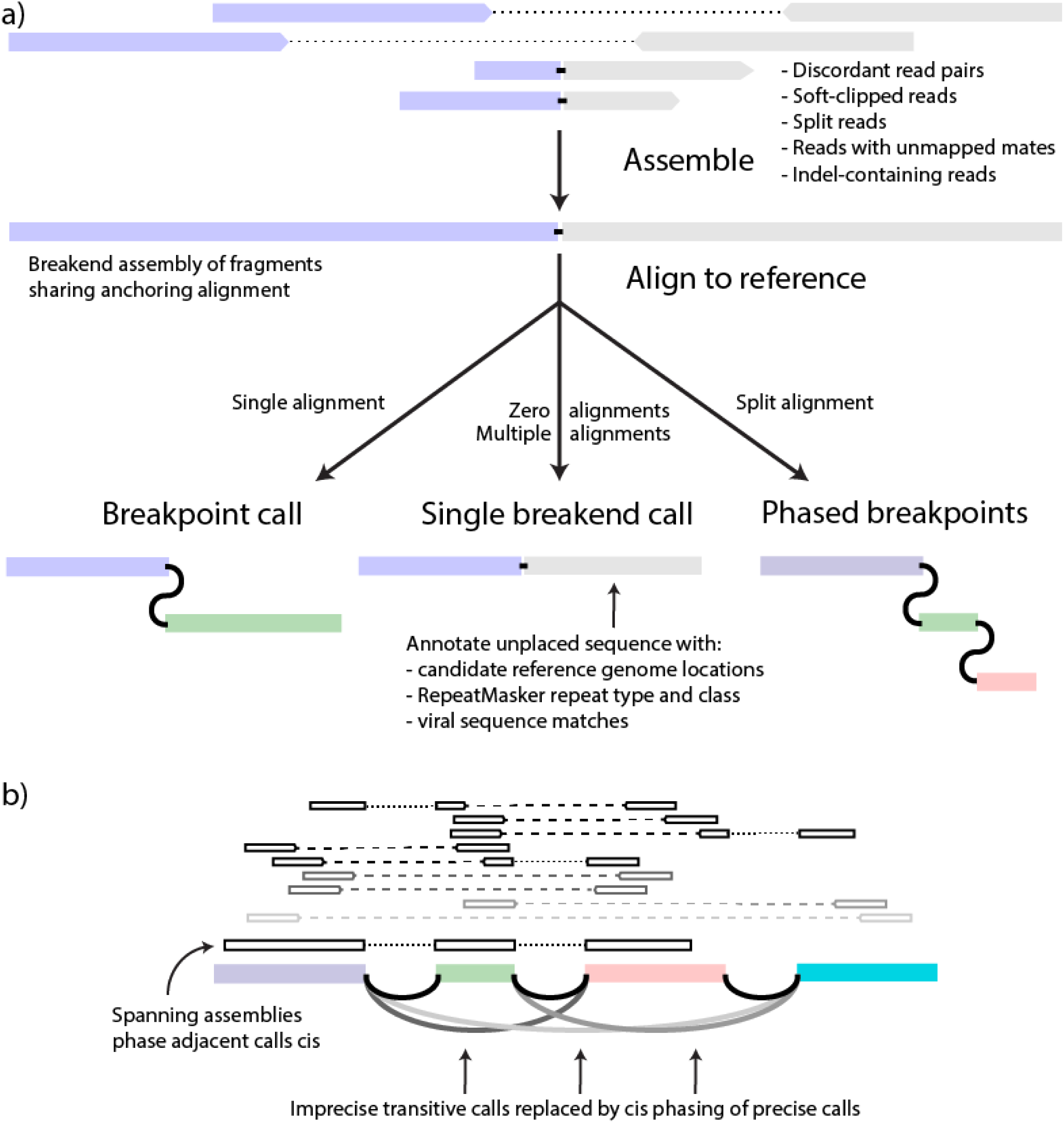
GRIDSS2 overview. a) contigs are assembled from a single locus of reads mutually supporting the same putative break junction. If the other side cannot be uniquely determined, the contig supports a single breakend call at the break junction position. If different portions of the contig sequence uniquely align to different genomic loci, the assembly supports multiple cis phased breakpoints. b) Nearby structural variants will have discordant read pairs spanning across multiple breakpoints. These generate spurious transitive calls that are collapsed into the underlying breakpoints, phasing them cis.

### Benchmarking performance

To estimate precision and sensitivity of GRIDSS2, we used a recently generated “gold standard” somatic SV truth set for the COLO829 melanoma cell line and the COLO829BL cell line, which was derived from a normal cell from the same individual, using a combination of Illumina, PacBio, Oxford Nanopore, 10X Genomics linked reads, and optical mapping followed by targeted capture and PCR-based validations and manual curation ^18^. To test sensitivity and reproducibility, we ran GRIDSS2, Manta^11^, svaba^19^, and novobreak^20^ on 3 independent sequencing replicates of the COLO829T/COLO829BL matched tumour-normal cell lines sequenced to a depth of 100x tumour and 40x normal coverage. GRIDSS2 achieved an average sensitivity/precision of 94%/83% compared to 88%/52% for Manta, 75%/11% for svaba and 70%/7% for novobreak (Figure 2a).

**Figure 2.**
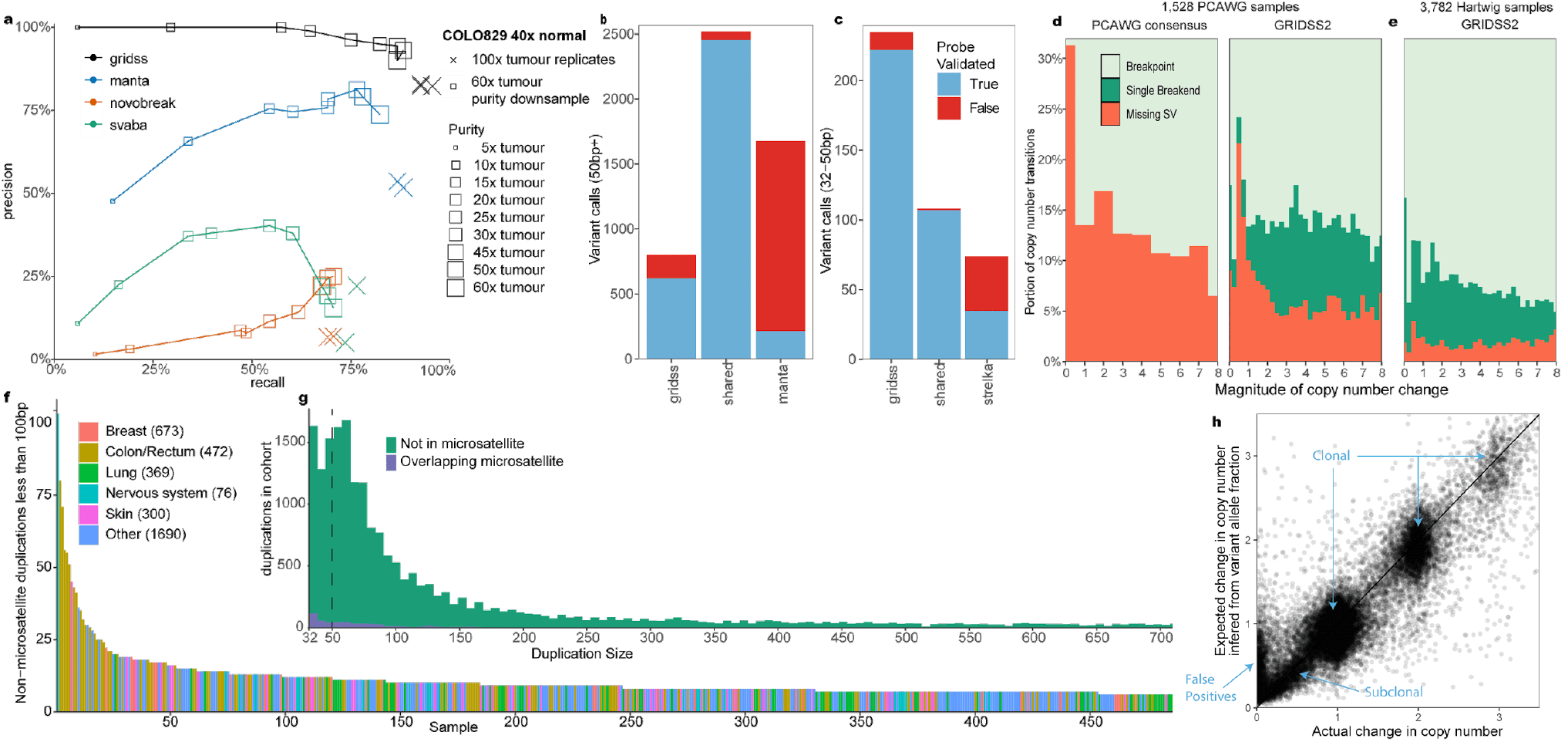
Somatic benchmarks. a) COLO829T/BL tumour and blood cell lines were sequenced in triplicate to 100x/40x. In-silico purity downsampling was performed at 40x normal, and 60x tumour coverage. Results are compared against a PCR validated somatic truth set generated from multiple sequencing technologies. b) GRIDSS2/Manta validation results on 13 patient samples for 50bp+ events. c) GRIDSS2/Strelka validation results for 32-50bp events. d) False negative rate (FNR) inferred from the presence of SVs copy number transitions broken down by magnitude of copy number change for 60 PCAWG samples. Comparison is between GRIDSS2/PURPLE and the PCAWG consensus call set. e) Inferred FNR for 3,782 100x tumour samples from the Hartwig cohort. Single breakend variant calling is crucial to the low FNR in this cohort. f) Per sample counts of 32-100bp somatic tandem duplications in the Hartwig cohort. These mutations are enriched in colorectal cancer and associated with ATM driver mutations. g) Size distribution of small (32-100bp) tandem duplications across the Hartwig cohort. This is a distinct signature not associated with microsatellite expansion. h) Comparison of expected vs actual copy number changes for the Hartwig cohort. SV inferred and actual copy number changes are closely correlated.

To evaluate performance at lower sequencing depths and sample purity, we use in-silico downsampling and mixing to simulate a matched normal at 40x and a 60x tumour sample at 8%-100% purity corresponding to 5x, 10x, 15x, 20x, 25x, 30x, 45x, 50x, and 60x effective tumour coverage. Above 10x effective tumour coverage GRIDSS2 achieved higher sensitivity and specificity than the benchmarked callers. At 10x and below, GRIDSS2 retained higher precision, but at lower sensitivity than Manta or svaba (Figure 2a).

### Validation on patient samples

To further validate somatic performance, we performed independent validation of GRIDSS2 and manta breakpoint calls from 13 patient tumor samples from the Hartwig cohort ^2,11,19,20^ with a high burden of structural variants. Since the default minimum reported event sizes of GRIDSS2 and Manta are 32 and 50bp respectively, we compared 32-50bp events to the short indel caller, Strelka ^21^. We used a hybrid capture approach with target probes flanking and overlapping break-junctions to independently validate over 5,000 calls identified by any tool. 3,403 of 3,666 (93%) GRIDSS2 calls were validated compared to 2,685 of 4,299 (65%) for Manta (Figure 2b). Of the private Manta calls not found by GRIDSS2, just 230 of 1777 (13%) were validated compared to 836 of 1031 (81%) GRIDSS2 private calls. Imprecise (that is, not base-pair accurate) Manta calls validated at a rate (40/288, 14%) similar to Manta private calls, whereas GRIDSS2 reports only precise somatic SV. No imprecise GRIDSS2 calls passed somatic filtering, whereas All validated imprecise Manta calls were called by GRIDSS2 precisely. In the 32-50bp range, 329 of 343 (96%) of GRIDSS2 calls validated against 142 of 182 (78%) for Strelka (Figure 2c). 95% (219 of 232) of 32-50bp calls private to GRIDSS2 were validated, compared to 47% (35 of 74) for Strelka. Notably, GRIDSS2 finds many short duplications of 32-100 bases which are largely missed by both Strelka and Manta.

### Novel somatic short duplication signature

In addition to reidentifying known kilobase and megabase length duplication signatures, we find a signature consisting of short 32-100bp non-microsatellite tandem duplications (Figure 2f). There is a median of 4 of these short (32-100bp) duplications per sample (Supplementary Figure 1). They are not correlated with larger duplications (R=0.08), or total breakpoints (R=0.10). Enrichment of samples with 15 or more short duplications is positively associated with colorectal cancer (Figure 2g) (q=1.2× 10^−9^) and driver mutations in PARK2 (q=0.0003) and ATM (q=0.008). Across the Hartwig cohort, 23 samples have driver mutations involving the disruption of a tumour suppressor caused by small duplications.

These short tandem duplications are too large to be reliably called by most somatic indel callers, but too short to be reliably called by many SV callers. In part this is due to the weak read pair signal due to the short variant length, but also since most callers do not report variants shorter than 50bp threshold used for variant databases such as dbVar. Popular callers such as lumpy ^22^ and delly ^23^ do not call duplications shorter than 100 and 300bp respectively ^4^, and no duplications shorter than 300bp were included in the PCAWG consensus call set^1^.

### Cohort-level FNR/FDR estimation using copy number consistency

Structural variant and copy number calls are intrinsically related. Any breakpoint must have either a compensating breakpoint (for example, as with inversions), or a copy number change at that SV position. Using this principle, we can estimate a false negative rate (FNR) from the number of unexplained copy number transitions. To generate matching SV and copy number calls, we ran GRIDSS2 and PURPLE ^2^ on 1,528 samples from the PCAWG WGS cohort and compared results with the state-of-the-art PCAWG consensus call set^24^. Copy number transitions in or within 100kb of centromeres or a gap in the reference genome were excluded.

Across the 1,528 samples, GRIDSS2 identified breakpoints for 84% of copy number transitions and single breakends for a further 4.7%, with an estimated 11.2% FNR. The PCAWG consensus call set identified breakpoints for 72% of copy number transitions (28% FNR). When restricted to clonal copy number transitions, the estimated FNR for the PCAWG consensus dropped to 14.2% and GRIDSS2 to 9.36% (Figure 2d), indicating robust subclonal GRIDSS2 performance.

To evaluate GRIDSS2 on high quality, deeply sequenced samples, GRIDSS2 and PURPLE were run on 3,782 40x normal/100x tumour samples from the Hartwig cohort. Excluding those occurring within 1kb of a gap in the reference genome, 153,231 of 1,954,548 (7.0%) copy number transitions in the Hartwig cohort were explained only by single breakend variants and 68,171 (3.1%) lacked a corresponding GRIDSS SV (Figure 2e). The higher rate of single breakend calling can be attributed to GRIDSS2 conservatively calling single breakends and the greater sequencing depth in the Hartwig cohort. The 7.0% of copy number transitions in the Hartwig cohort explained by single breakend variant calls represents a lower bound for the FNR of an exclusively breakpoint-based caller. A FNR of 3.1% suggests that, on this cohort, GRIDSS2 achieves a FNR lower than that possible for a breakpoint-based caller.

To demonstrate that this reduction in FNR does not come at the cost of a high false discovery rate (FDR), we compared the change in copy number to the change expected based on the variant allele fraction (VAF). For isolated breaks, the change in copy number should match the variant copy number inferred from the variant allele fraction. Using a 3000bp threshold to ensure at least one full 1kbp copy number bin between SVs, we find that the VAF-inferred SV copy numbers reported by GRIDSS2 are consistent with the copy number changes with no systematic bias in the VAF (Figure 2h). This trend remains true for subclonal variants although the false discovery rate does go up. Assuming variants with a copy number change of less than 0.1 and a VAF inferred copy number of at least 0.25 are false positives, GRIDSS2 isolated SV calls have an estimated FDR of 5.4%, with 74% of these subclonal, and single breakends having twice the FDR of breakpoints. Extrapolating these to the rest of the cohort gives an overall estimated FDR of 3.3%.

### Resolving somatic centromeric rearrangements

Although only one side of single breakend variant calls can be uniquely placed, the assembled sequencing flanking the break can be used to classify integrated provirus, mobile element transposition, rearrangements involving centromeric and telomeric sequence, and other events. RepeatMasker annotation reveals that the majority of somatic single breakend calls are caused by SINE Alu, LINE L1HS insertions or rearrangements involving centromeric sequence, a pattern shared between both the Hartwig and PCAWG cohorts (Figure 3a, Supplementary Figure 2). Breakend assembly lengths for SINE single breakends are typically shorter than 150bp as assemblies longer than this can typically be resolved into breakpoint calls. Similarly, the polyA repeat motif characteristic of LINE translocations^25^ is also found in the shorter breakend assemblies. Such assemblies are short as the de Bruijn graph assembler used truncates assemblies at unresolved repeat loops and assemblies able to span the polyA tail are able to be resolved as breakpoints.

**Figure 3.**
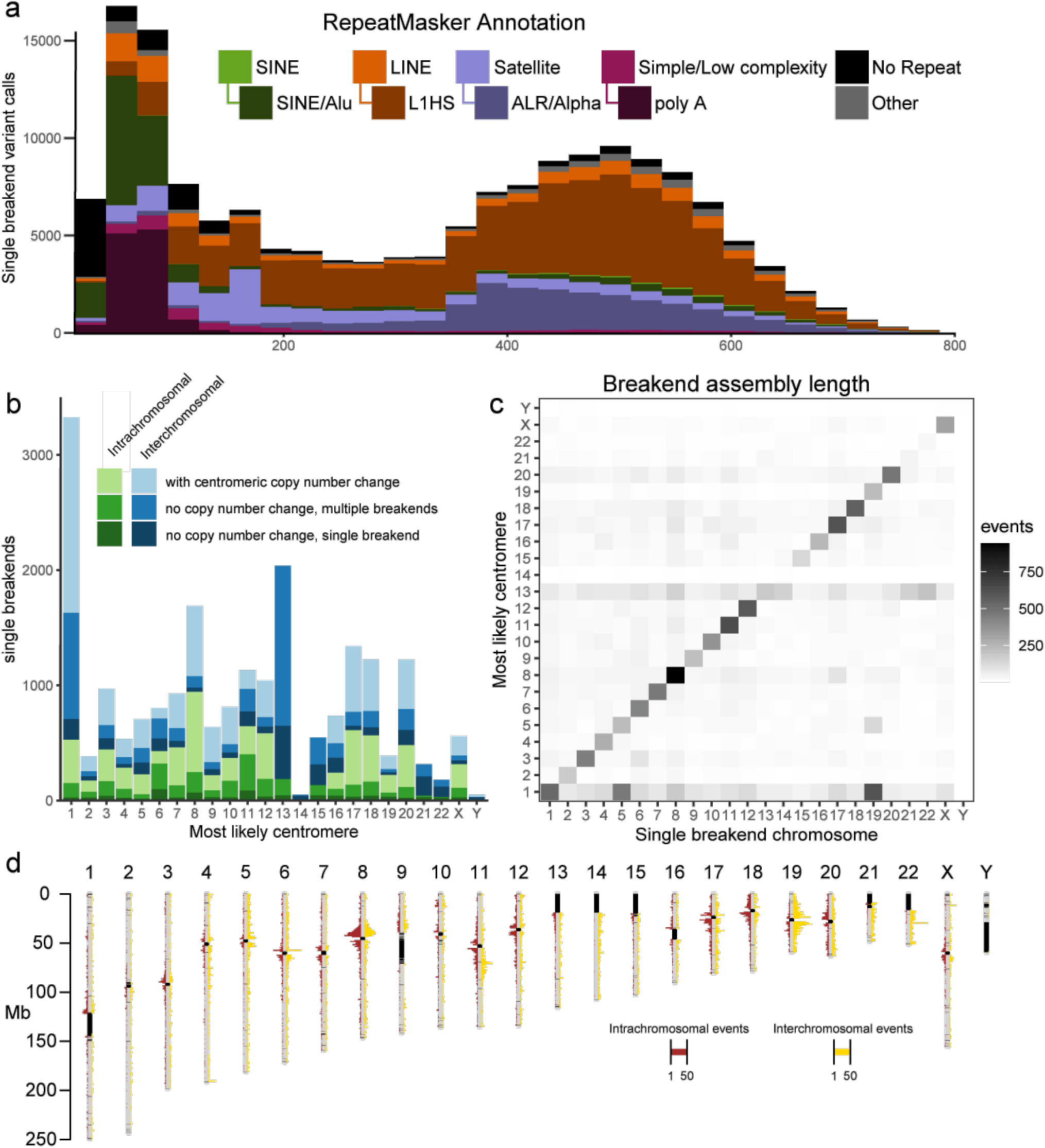
Classification of single breakends. a) RepeatMasker annotations indicate the majority of somatic single breakends are due to mobile element translocations, or centromeric breaks. b) Most likely centromere for single breakends containing centromeric or peri-centromeric repeats based on realignment of breakend sequence to hg38. Shading indicates whether prediction is consistent with the copy number change across the centromere. Chromosome 1 has an excess of inter-chromosomal breaks to centromeric sequence. Chromosomes 13, 14, 15, 21, 22 have insufficient non-gap p-arm sequence for a centomeric copy number change to be called. c) Location of single breakends to centromeric sequence and corresponding centromere. Chromosome 1 has an excess of inter-chromosomal breaks to centromeric sequence, particularly to 5 and 19. d) Location of single breakends connected to centromeric sequence on the same chromosome. Red events left of the chromosomes are intra-chromosomal, and yellow events to the right are inter-chromosomal.

91% of the Hartwig cohort samples contain at least one copy number transition occurring in centromeric sequence. Being able to resolve the partners of the centromeric breaks explaining these copy number changes is critical to the accurate reconstruction of the derivative chromosomes. Single breakends into ALR/Alpha and HSATII centromeric repeats are able to give significant insight into the nature of these centromeric breaks. As each human centromere has a slightly different dominant repeat sequence, a mapping between each centromeric single breakend and their most likely centromeric breakpoint partner is possible. To do this, we aligned the single breakend sequences containing a centromeric or peri-centromeric repeat against the hg38 reference genome using BLAT, annotating each with the most likely centromeric partner. Using this approach, we were able to explain 5,614 of 11,996 (47%) centromeric copy number changes, implying that approximately half of centromeric rearrangements are centromere to centromere, and the remainder centromere to non-centromeric sequence. Of the 21,587 centromeric single breakends detected 3,148 (15%) had no copy centromeric copy number change, 6,850 (32%) had no copy number change but had multiple single breakends linked to the same chromosome, 3,358 (16%) had a single breakend associated with a centromere with copy number change, and the remaining 8,231 (38%) associated with a centromere with copy number change with multiple breakends mapping to that centromere in that sample.

### Novel centromeric break signature

The centromeric single breakend rate can be further broken down by chromosome (Figure 3b) and based on the location of the single breakend (Figure 3c). Chromosome 1 is a clear outlier with an overabundance of centromeric inter-chromosomal rearrangements, particularly to chromosomes 5 and 19. Although the high level of sequence similarity between the centromeres of 1, 5, and 19 ^26^ could be a cause of false positive predictions, this relationship holds even when restricting the analysis to single breakends with an associated centromeric copy number change (Supplementary Figure 3), implying that the centromeric similarity between 1, 5 and 19 results in an increased rate of centromeric rearrangements between these chromosomes. In contrast, the lack of copy number supported single breakends to chromosome 13,14, 15, 21, and 22 centromeres is an artifact caused by missing p arm copy number due to gaps in the reference genome. Similarly, the centromeric sequence homologies between 13, 14, 21, and 22 combined with the lack of confirmatory copy number support, make it difficult to determine how much of the high inter-chromosomal centromeric rearrangement of chromosome 13 is due to misattribution of rearrangements to other chromosomes.

In general, intra-chromosomal single breakends to centromeric sequences occur close to the centromere (Figure 3d), with this effect less pronounced for inter-chromosomal breaks. Chromosome 1 is enriched for inter-chromosomal breaks, particularly to chromosomes 5 and 19, with inter-chromosomal breaks from these chromosomes to the centromere on 1 (Supplementary Figure 4) occurring in a pattern similar to the intra-chromosomal breaks of other chromosomes.

### Somatic phasing

The breakend assembly approach taken by GRIDSS2 also enables the assembly-based phasing of nearby variants. When two structural variants occur in close proximity, they can be phased as cis if the contig aligns across both, and trans if the contig aligning across one aligns to the reference sequence at the other (Figure 4a). Segments shorter than 30bp are not typically uniquely alignable by BWA and unaligned short DNA segments are treated as insert sequences of an SV connecting the longer flanking segments. Since breakend assembly contig lengths are limited by the fragment size distribution of the DNA library sequenced, only nearby variants can be phased.

**Figure 4.**
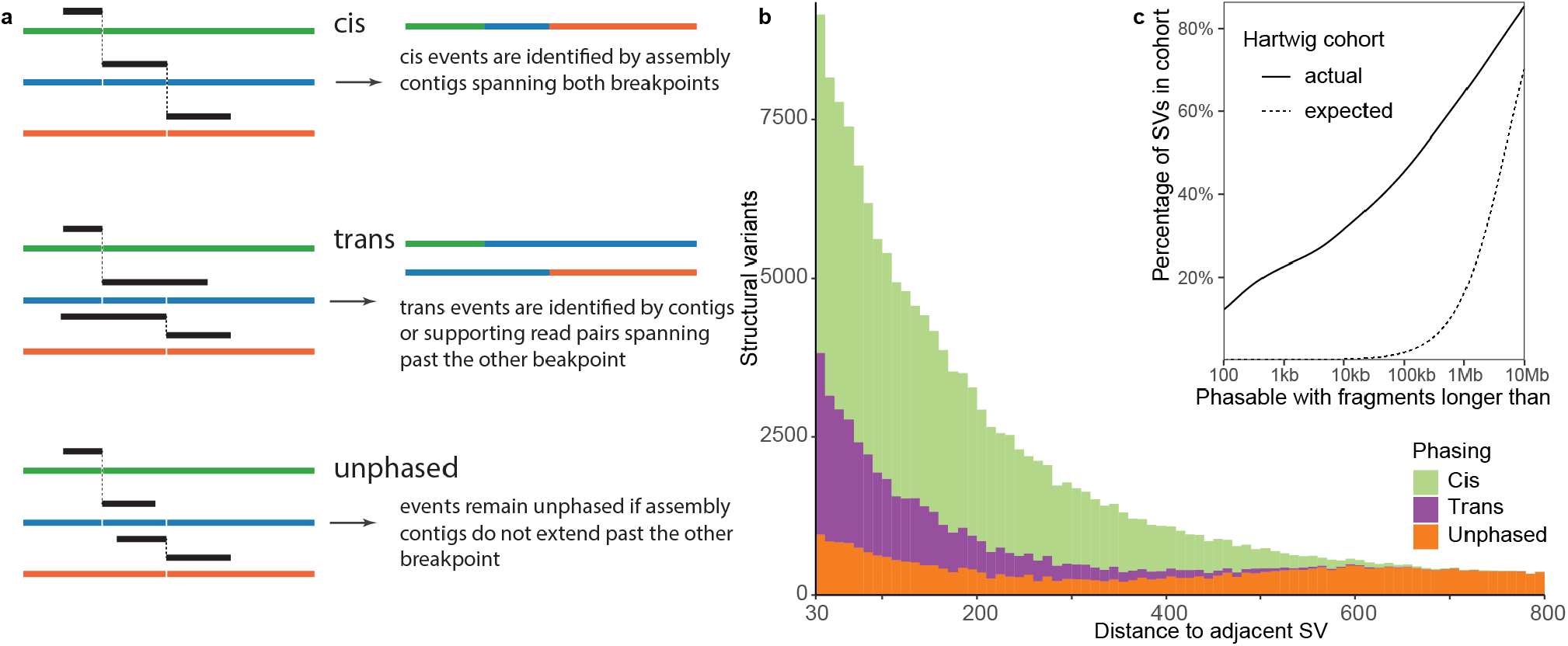
Structural variant phasing. a) Phasing of structural variants can be determined when breakend assembly contigs span multiple breakpoints. b) The majority of variants within 600bp can be phased using breakend assembly. c) Somatic SVs are highly clustered with 22% of all SVs in the Hartwig cohort potentially involving a DNA fragment of 1kbp or less.

For the Hartwig cohort, variants could be phased up to around 500 base pairs. We found that multiple nearby somatic structural variants are frequent, with 22% of all structural variants having an adjacent variant within 1,000bp. This is far in excess of the 0.02% expected if the breakpoints were uniformly randomly distributed (Figure 4c). Of these, GRIDSS2 was able to phase 70% (Figure 4b) with 72% cis and 28% trans. This distribution is recapitulated in the 1,528 PCAWG samples and LINX classification of these structural variants indicate that that phasable breakpoint clusters occur predominantly in LINE translocations (due to target site duplication, and highly active donor elements) and highly complex rearrangement events (Supplementary Figure 5). This phasing information greatly assists downstream derivative chromosome reconstruction, as it exponentially reduces the number of possible paths through the breakpoint graph.

### Impact on complex event resolution

To demonstrate the impact on downstream analysis of complex somatic genomic rearrangements, we ran LINX ^27^, a rearrangement event interpretation and classification tool, on the Hartwig and PCAWG cohorts. To fully resolve complex rearrangements structural variants must be chained together to reconstruct the relevant portions of the derivative chromosomes rearranged by the event. If there are errors in the SV call set, it is likely that many complex events will not be able to be fully resolved. To evaluate the impact of FNR on this reconstruction, we evaluated the portion of SVs resolved into long chains for the PCAWG and Hartwig cohorts. In addition, we simulated the effect of increasing FNR by subsampling the Hartwig call set (Figure 5a). 5.0% of SVs in the Hartwig cohort were reconstructed into chains of 20 SVs or more. Increasing the FNR reduces this to 3.6% of SVs at 5% FNR, 1.5% at 10% FNR, 0.6% at 15% FNR, and 0.25% at 20% FNR. We previously estimated the PCAWG cohort FNR at 11.2% and we find that the 1.27% of SVs in chains of 20 SVs or more closely match the 1.29% we expect from a simulated downsampling of the Hartwig cohort. This implies that the PCAWG and Hartwig pancaner cohorts have a broadly similar composition of complex rearrangements and the differences observed are primarily technical in nature. Small improvements in FNR result in large increases in the ability for downstream tools to resolve complex events. A sub-5% FNR is critical for large event reconstruction.

**Figure 5.**
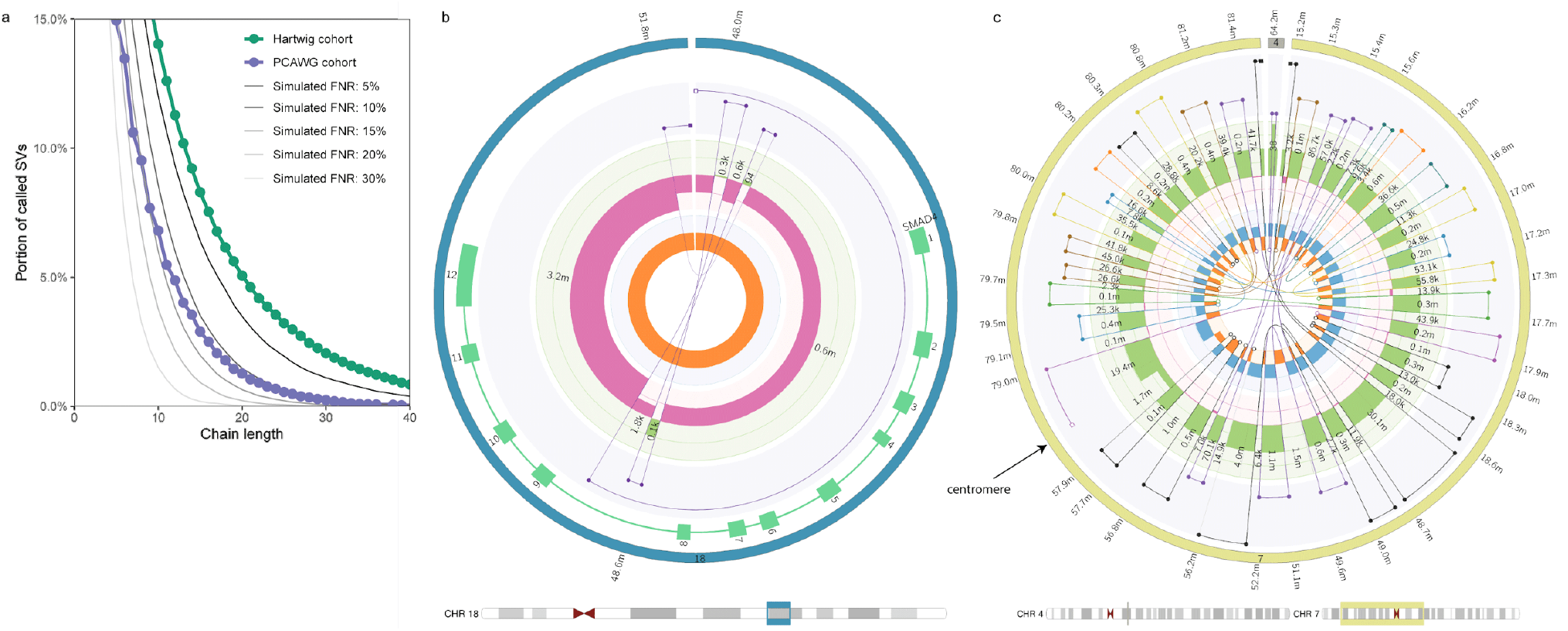
Complex rearrangement interpretation. a) Impact of false negative rate (FNR) on complex event resolution. The y-axis indicates the portion of structural variants that form part of a resolved chain of SVs at least as long as the chain length indicated on the x-axis. LINX results for GRIDSS2 calls on the Hartwig and PCAWG cohorts are shown along with simulated results from downsampling the Hartwig cohort to the specified FNRs. A low FNR is essential to accurate complex event resolution. b) Circos plot of SMAD4 driver deletion event. The interpretation of this deletion is confounded by the presence of 3 short fragments at the breakpoint. This event can be fully resolved by GRIDSS2 SV phasing. Circos tracks from innermost to outermost are: single breakends (open white circles) and breakpoints, LOH, copy number, connected segments, genes, chromosome. b) Circos plot of chromothripsis overlapping centromeric sequence. This event spans across the chromosome 7 centromere. A subset of the chromothriptic fragments have been inserted into chromosome 4. Each SV chain is represented in a different colour.

SV phasing can be critical to the correct interpretation of complex events. For LINX, SV phasing is a critical first step in the chaining of SVs. Of the 486,632 links in the chains resolved by LINX in the Hartwig cohort, 100,007 (21%) were due to GRIDSS2 SV phasing. For the PCAWG cohort, 13,212 of 107,952 links (12%) were resolved by GRIDSS SV phasing, with the difference primarily driven by shallower coverage and shorter library fragment sizes resulting in shorter assembly contig lengths (Supplementary Figure 2), and the higher FNR. In some cases, apparently complex events can be resolved to simple events containing additional short DNA fragments purely through SV phasing (Figure 5b).

Finally, we use single breakend repeat annotations to identify instances of chromothripsis overlapping centromeres. In the Hartwig cohort, LINX identifies 270 complex events with at least 10 breakends to centromeric sequence, 17 of which could be fully chained (Figure 5c). The large number of events with many centromeric single breakends indicates a previously unexplored level of centromeric involvement in complex rearrangements worthy of further investigation.

## Discussion

Through cell line, patient validation, and cohort-level comparisons, we have shown GRIDSS2 has excellent somatic performance above 10x effective tumour coverage. The identification of a small (32-100bp) duplication signature by GRIDSS2 highlights the importance of robust software tested across a wide range of variant types and sizes. The presence of a signature overlapping the widely accepted but arbitrary 50bp threshold separating indels from structural variants suggests it is time to reconsider this threshold as the minimum reported event size for structural variants.

Explicitly reporting and handling of single breakend variants represents a significant conceptual advancement in the treatment of genomic rearrangements. Even though only the high-mappability side can be unambiguously placed, sequence classification of the low-mappability side produces useful results and meaningful insights. The identification of frequent somatic centromeric rearrangements demonstrates the utility single breakend variant calling has in regions of the genome traditionally considered inaccessible to short read sequencing. Single breakend variant calling provides a framework for the reliable detection of LINE integration without a specialised caller ^28^, for the detection of centromeric and telomeric viral integrations ^29^, and for an entire ecosystem of tools that explicitly model the ambiguity they represent. As single breakends comprise 7.0% of GRIDSS2 calls in the Hartwig cohort, any purely breakpoint-based caller must have a false negative rate of at least 7.0%. GRIDSS2’s 3.1% FNR may thus be impossible to achieve for any breakpoint-based caller, at least for this cohort.

One biologically significant finding coming from GRIDSS2’s ability to phase structural variants is the degree to which somatic structural variants are clustered. In the Hartwig cohort of metastatic solid tumours, 22% of somatic structural variants potentially involve DNA fragments of less than 1000bp with GRIDSS2 able to phase 70% of these. Long read sequencing is considered the gold standard for structural variant phasing and phasability is indeed better: 10kb long reads increase this theoretical phasability to 31%. The high indel error rate of PacBio and ONT sequencing presents a current drawback of long read sequencing: simple long read based SV detection approaches are likely to misidentify complex rearrangements involving nearby cis-phased SVs. Without HiFi sequencing or error correction prior to alignment, the short DNA segments between the SVs will be unmappable and the long read caller will report a transitive call between the flanking segments as outlined in Figure 1b. On COLO829, we found 5 instances (of 67 true positives) in which GRIDSS2 based on short-read sequencing was able to correctly place a short DNA segment that the three long-read callers were not. Care must be taken when comparing or combining short and long read variant calls to ensure the different representations of the same event are reconciled and cis phased. GRIDSS2’s ability to phase breakpoints involving short DNA fragments is of great utility to downstream rearrangement event classification and karyotype reconstruction as it exponentially reduces the number of possible paths through the breakpoint graph. The highly clustered nature of somatic SVs means that short read sequencing is surprisingly competitive when it comes to phasing somatic variants. Sophisticated analysis and interpretation of somatic genomic rearrangements does not necessarily require long read sequencing.

Single breakend variant calling enables a sensitivity and specificity unprecedented amongst short read-based somatic structural variant callers, facilitating the resolution of highly complex rearrangements. While breakpoints and copy number segments are widely adopted fundamental genomic rearrangement signals, single breakends have been hitherto unutilised. Their introduction enables the ambiguities present in low mappability regions to be explicitly modelled without compromising FNR or FDR and their potential extends far beyond the examples presented here. GRIDSS2 demonstrates that single breakend variant calling is essential to the comprehensive characterisation of somatic structural variation from short read sequencing data. Combining single breakend variant calling and structural variant phasing with low FNR and FDR, GRIDSS2 represents a foundation upon which sophisticated somatic analysis can be performed.

## Methods

GRIDSS2 extends the GRIDSS^12^ software suite with additional features, tools and capabilities. GRIDSS2 is composed of the following 5 phases: (i) preprocessing, (ii) assembly, (iii) variant calling, (iv) annotation, and (v) somatic filtering.

### Preprocessing

GRIDSS2 takes one or more aligned samples in the SAM/BAM^30^ file format. These files are pre-processed on a per-file basis and all reads supporting a structural variant are extracted, and all fields or tags referring to another record are corrected. Reads with chimeric alignments (i.e. split reads), reads with a soft or hard clipped alignment CIGAR of at least 5bp, read pairs in which only one read is mapped, and discordant read pairs are extracted. The library insert size distribution is estimated from the first 10,000,000 reads using picard tools (http://broadinstitute.github.io/picard) and read pairs considered discordant if they fall outside the 99.5% distribution of fragment size lengths. The clipped bases of soft clipped non-chimeric reads are realigned to the reference genome using bwa mem ^31^ and converted to a chimeric split read alignments if an alignment is found. Inconsistencies in the mate chromosome and position are corrected (since tools such as GATK indel realignment adjust read alignment positions without updating the mate record), hard clips converted to soft clips, the NM, SA, MC, MQ tags recalculated, and the R2 tag is populated. Improving performance over GRIDSS, GRIDSS2 performs this in a two-pass manner with samtools^30^ used for name/coordinate sorting the output of the first/second pass respectively.

As with GRIDSS, reads with low alignment sequence entropy and reads with a mapping quality (mapq) less than 20 (c.f. mapq<10 GRIDSS) are treated as unmapped, soft-clipped reads with clipped sequence having high homology with standard adapter sequences are ignored, reads marked as duplicates, and regions above 50,000x (c.f. 10,000x) coverage are ignored. Read alignments containing an insertion or deletion under 5bp are considered consistent with the reference.

### Assembly

GRIDSS2 uses the same genome-wide positional de Bruijn graph break-end assembler used by GRIDSS. Reads are split into kmers and associated positions based on the anchoring alignment: kmers from split reads must be assembled only with kmers at the positions inferred by the anchoring alignment, and kmers of unmapped mate reads are assembled at any position consistent with the library fragment size distribution and the anchoring read alignment position. For assembly purpose, split reads and indel alignments are considered multiple soft clipped alignments, and discordant read pairs are treated as multiple read pairs with one read aligned.

The output of the assembly is a set of ‘soft-clipped’ contigs with anchoring bases supporting the reference, and non-reference bases supporting a putative breakpoint at a given position. This contig is iteratively realigned to the reference using bwa mem and converted to a split read alignment. Assemblies longer than the 1.5x maximum fragment size distribution, as well as assemblies supporting the reference sequence are ignored. Assembly alignments with a mapq of less than 20 are treated as unmapped. GRIDSS2 introduces a number of refinements to the assembly calling process.

Assembly support is tracked per base pair. Fragments are considered to support a breakpoint only if the fragment support spans at least one base pair beyond any breakpoint homology on both sides. This ensures that when a single contig spans both a germline indel and a somatic SV, the fragments originating from the matched normal sample will not be considered as supporting the somatic breakpoint. This also improves variant allele fraction calculations in regions of complex rearrangement.

GRIDSS2 performs compound realignment of the entire assembly contig. BWA is used to align the entire assembly contig. Assembly contig bases which are soft clipped in the primary alignment reported by BWA are fed back to BWA for realignment. This process is repeated until either all bases are aligned, or no alignment can be found for the remaining bases. Assembly contigs that do not overlap with the locus of origin of the assembly are filtered out. To ensure that valid assemblies are not unnecessarily filtered, GRIDSS 2 includes both reads of each fragment in the assembly, and up to 300bp of anchoring reference-supporting sequence is included in the assembled contig. The remaining contigs are treated as split read alignments. To rectify over-alignment of the primary alignment location in the presence of imperfect breakpoint microhomology, the bounds of each split are adjusted to minimise the edit distance to the reference. The originating alignment is tracked using OA SAM tag and contigs that do not partially align to the originating assembly location and strand are filtered.

gridss.SoftClippedToSplitReads invokes bwa, -L 0,0 is added to the command line to remove the soft-clipping alignment penalty. This prevents 1bp non-template inserted sequences being over-aligned and reported as clean breakpoints with a flanking SNV.

Worse-case assembly performance has been improved by adding an assembly graph path count threshold. Generating 3 assembly contigs with more than 50,000 alternative paths through the assembly graph without advancing the assembly window will flush the assembly window. The maximum assembly window size has been reduced by 2.5x and more aggressive assembly read downsampling in high coverage regions is performed.

The presence of a contig with at least three non-overlapping alignments results in the breakpoints supported by that assembly being phased cis. If the initially soft-clipped portion of an assembly realigns across one breakpoint but not another, these breakpoints are phased trans.

### Variant Calling

Breakpoints are called using a probabilistic model based on the empirical distribution of CIGAR operators, the library fragment size distribution, and mapping rate. Each read/read pair is given a phred-scaled quality score based on the mapping quality and the probability of encountering that read/read pair given the library empirical distribution. Split reads and soft clipped reads use the distribution of soft clipping CIGAR operators. Discordant read pairs use the discordant mapping mate if distal or the library fragment size distribution if falling within the range reported by Picard tools CollectInsertSizeMetrics. Reads with unmapped mates use the unmapped mate fragment mapping rate, and indels based on rate of alignments with insertion/deletion CIGAR elements of matching lengths. As with GRIDSS, split reads and breakpoint-supporting assemblies incorporate the mapping quality scores on both sides of the supported break.

The key novel feature of the GRIDSS2 variant calling processing is the reporting of single breakend variants. Single breakends variants are called based on supporting soft clipped reads, assembly contigs, and reads with unmapped mates. Single breakend calling uses the same two-pass approach as breakpoint calling with all maximum cliques first calculated, then evidence uniquely assigned to the highest scoring clique.

In addition to single breakend variant calling, the variant caller has been improved by: reducing the default minimum called event to 10bp; preferentially allocating reads to variants supported by an assembly containing the read; requiring two supporting fragments to call a variant; and excluding inversion-like breakpoints from the minimum variant size filter to prevent filtering of foldback inversions.

### Annotation

GRIDSS2 provides a full per-sample breakdown of all supporting evidence for each variant through the following VCF INFO and FORMAT fields:

- AS, RAS, CAS: counts of assembly contigs supporting a breakpoint originating locally, from the other side of the breakpoint, and from another location respectively. CAS assemblies support multiple variants and provide linking information about those variants.
- ASSR, ASRP: total number of split/soft clipped/indel-containing reads, and discordant read pairs/reads with unmapped mate contributing to any breakpoint-supporting assembly contig at the breakpoint location. Note that read/read pairs that are assembled into a contig but whose interval of support does not span the breakpoint are not counted. The interval of support for a read/read pair is defined as the interval between the first and the last contig base for which that read/read pair contributed to the assembly.
- SR, RP, IC: counts of split reads and discordantly mapped read pairs, and indel-containing reads that directly support the breakpoint.
- BA: counts of assembly contigs support a single breakend at this position. Such contigs are aligned only to the local breakend with the breakend sequence either aligning ambiguously, or unable to be aligned to the reference genome by bwa.
- BASSR, BASRP: total number of split reads or soft clipped reads, and discordant read pairs or reads contributing to any breakend-supporting assembly contig at the variant location.
- BSC, BUM: counts of soft-clipped reads, and reads with unmapped mates at the variant location
- ASQ, RASQ, CASQ, SRQ, RPQ, IQ, BAQ, BSCQ, BUMQ: corresponding quality score contribution for the supporting evidence.
- QUAL, BQ: total contribution to the breakpoint/breakend quality score.
- BANRP, BANSR, BANRPQ, BANSRQ: counts of read pairs/split reads not supporting this breakpoint but assembled into a contig that supports this breakpoint and their corresponding assembly quality score contribution.
- REF/REFPAIR: count of reads/read pairs spanning the local variant position that support the reference allele. Only reads/read pairs that span across the breakpoint microhomology interval (if present) are counted.
- VF/BVF: count of unique fragments supporting the breakpoint/breakend. By tracking unique fragments supporting the variant, a more accurate variant allele fraction can be calculated. This approach prevents double-counting of discordantly mapped fragments for which one of the reads contains a split read alignment. A fragment can support a variant either directly through split read, soft clipped read or discordant alignment of a read pair, indirectly through incorporation of one or both of the constituent reads in an assembly supporting the variant, or both directly and indirectly.
- RF: count of unique fragments supporting the reference allele.
- CQ: variant quality score prior to evidence reallocation.
- BEALN: Potential alignment locations of breakend sequence as determined by *gridss*.*AnnotateInsertedSequence*.
- BEID, BEIDL, BEIDH: identifiers of assembly contigs and the corresponding local and remote alignment base offsets. Single breakend variants do not have a remote breakend, and only breakpoint variants include breakpoint-supporting assemblies. Variants containing the same BEID are phased cis.
- CIPOS: For IMPRECISE variants, CIPOS encodes the interval in which the breakpoint could occur and for precise variants, CIPOS encodes the homology interval.
- CIRPOS: corresponding CIPOS of the remote breakend.
- IHOMPOS: interval of inexact homology. A Smith-Waterman alignment of the breakpoint sequence against the reference sequence is performed at both breakends. The reference and breakpoint sequence are extended 300bp from the break on either side with the reference extended an additional 10bp to account for potential indels in the alignment. The homology length is the length that the sequence alignment could be extended from the common sequence into the breakpoint/reference sequence. Alignments containing a soft clip on the common sequence side are classified as alignment errors and ignored. The SSW library^32^ is used for which we implemented a JNI wrapper. Alignment scored 1, -4, 6, 1 for match, mismatch, gap open, and gap extend respectively which correspond to bwa mem alignment scores.
- SC: CIGAR encoding of the anchoring bases that at least one read/read pair/assembly is aligned to and supports the variant. This is encoded as a CIGAR string with a match for each anchoring base that provides support for the variant call, XNX for the interval over which the breakpoint could occur (due to microhomology or an imprecise call), and a deletion CIGAR element for any intervals over which there is no support (such as a small flanking deletion). Variants with an anchoring SC 10bp further from the break than a nearby variant are considered to be phased trans.
- SB: Strand bias of the reads supporting the variant. 1 indicates that reads would be aligned to the positive strand if the reference was changed to the variant allele. 0 indicates that reads
- bases would be aligned to the negative strand if the reference was changed to the variant allele. Strand bias is calculated purely from supporting reads and exclude read pair support since these are intrinsically 100% strand bias. Note that reads both directly supporting the variant and supporting via assembly will be double-counted. Both breakpoint and breakend supporting reads are included.
- IMPRECISE, HOMLEN, HOMSEQ, PARID, EVENT, CIEND, END, and SVTYPE fields carry their usual meaning as per the VCF file format specifications.
- MQ, MQN, MQX, BMQ, BMQN, BMQX mean, min, and max MAPQ score of reads/assembly contigs providing breakpoint/breakend support.

After initial annotation, *gridss*.*AnnotateInsertedSequence* aligns any single breakend sequences or non-template inserted breakpoint sequences to an arbitrary reference genome and adds an annotation reporting the alignment location. Integrated viral sequence is identified by aligning to a reference of viral sequences. By default, the same reference as the input files were aligned to is used. If a RepeatMasker bed file generated by BedOps^33^ rmsk2bed is supplied, inserted sequences will be annotated with the RepeatMasker class and type corresponding to the BEALN alignments.

### Somatic filtering

By default, GRIDSS2 is a sensitive caller and reports all putative variants supported by at least two well-mapped reads. To generate a set of high and low confidence somatic call sets, a somatic filter was developed. Variants with 3% of the supporting reads originating from the normal, or deletion or duplication breakpoints under 1000bp that have any direct split read support in the normal, are hard filtered. Variants are classified as low confidence if any of the following conditions are met: breakend coverage of less than 8 fragments in the normal; allelic fraction of less than 0.5% in the tumour; imprecise variant call; breakend variants without an assembly containing at least one discordant read pair; single breakends with a poly-C or poly-G run of at least 16bp in the breakend sequence; deletion or duplication breakpoints under 1000bp with a split read strand bias of 0.95 or greater; breakpoints with a microhomology of over 50bp; breakpoints with an inexact microhomology of over 50bp which are not deletion or duplications under 1000bp; deletion or duplication breakpoints under 1000bp with no split read support either directly, or through assembly; breakpoints with no discordant read pair support (either directly, or via assembly) which are not deletions or duplications under 1000bp; deletion or duplication breakpoints under 1000bp that have any direct split read support in the normal; 100-800bp deletion breakpoints with an inexact microhomology length of 6bp or greater; inversion-like breakpoints 40bp or less that have at least 6bp of microhomology; deletion-like breakpoints under 1000bp whose length of sequence inserted at the breakpoint is within 5bp of the deletion length, except those whose edit distance to the deleted bases is at least 0.5 per base, and less than 0.2 per base to the reverse complement. Breakpoint variants are filtered if either breakend is filtered.

Somatic variants are panel-of-normal (PON) filtered if a match within 2bp was found in a panel of normals. The default hg19 was constructed from the 40x coverage WGS matched normals for 3,972 patients from the Hartwig cohort using the *gridss*.*GeneratePonBedpe* utility. If multiple samples for a patient existed, only the normal for the first sample was included in the PON. Variants were aggregated across samples using the default setting of ignoring the FILTER field, and excluding imprecise calls and breakpoints/single breakends with a QUAL score of less than 75/428.

Viral insertions are annotated using *gridss*.*AnnotateInsertedSequence*. Single breakend sequences and non-template inserted sequences that do not have an alignment to the reference genome were aligned to a set of human viral reference sequences. Viral reference sequences were obtained from the virus host database ^34^ and filtered to include only viruses associated with the homo sapiens taxid of 9606. The viral sequences were then masked using RepeatMasker with “-no_is -s -noint -norna -species human” parameters. Generation scripts can be found at https://github.com/hartwigmedical/scripts/tree/master/virus.

Assembly linking: pairs of breakpoints mutually supported by a common assembly contig were annotated as linked by assembly. For assembly contigs spanning more than 2 breakpoints, each adjacent pair was linked with a unique identifier to enable unambiguous traversal of the breakpoint graph.

Transitive linking: chains of precise breakpoint variants were phased trans if an imprecise spanning transitive breakpoint call could be found. To identify transitive calls, a breadth-first search over the breakpoint graph was performed. Variants were considered transitive if the start and end breakends overlapped the start and end breakpoints in a path of precise breakpoint calls. Paths were limited to 1,000bp and 4 segments, with each segment required to be at least 20bp in length. Paths could not self-intersect. To prevent exponential runtime in highly rearranged genomes, at most 100,000 paths and at most 1,000 paths per starting breakpoint were considered.

Simple inversion annotation: pairs of breakpoints with orientations consistent with a simple inversion were annotated as simple inversions if the matching breakends were within 35bp on both sides, no other simple event annotation could be applied, and fragments supporting the constituent variants differed by at most threefold.

Templated insertion annotation: breakend/breakpoint and breakend/breakend pairs were annotated as simple templated insertions if the breakends had opposite orientations, were within 35bp, no other simple event annotation could be applied, and fragments supporting the constituent variants differed by at most threefold.

Reciprocal translocation: breakpoint/breakpoint pairs were annotated as reciprocal translocations if the breakends on both sides had opposite orientations, were within 35bp, no other simple event annotation could be applied, and fragments supporting the constituent variants differed by at most threefold.

Equivalent: variants were annotated as equivalent if variants had a breakend within 5bp of each other and they shared a common breakend sequence. Breakend sequences were truncated to the length of the shorter sequence and were considered matching when the per-base edit distance between breakend sequences was 0.1 or less. For the purposes of this comparison, the nominal breakend sequence was used for single breakends, and the reference sequence of the partner breakend was used for breakpoint variants. For breakpoint variants, the length of the breakend sequence was the maximum of 20 bases, and the width of the interval over which the fragments supporting the partner breakend had anchoring alignments.

Finally, a quality filter was applied to breakpoint variants with a QUAL score of less than 350 and single breakend variants with a QUAL score of less than 1000. Variants linked to a variant passing the qual filter other than through equivalence were rescued from the quality filtering and were considered to have passed regardless of the actual variant quality score. For each input file, two output files were generated: a high confidence call set containing calls passing all filters and a low confidence call set containing all calls except those failing the normal support filter or short events with split read support in the normal.

### Independent validation of SV calls

13 samples from the Hartwig metastatic cancer cohort were selected for capture panel validation of the structural variant calls. Each variant called in GRIDSS2 was compared with variants called from Manta (for variants longer than 50 bases) and/or Strelka (for variants from 32-50 bases in length) to determine if the variant is shared or private. Variants were marked as matching GRIDSS2 if the start and end chromosomes and orientation both matched and start end positions (including confidence intervals) were within 20 bases of each other.Hybrid capture probes were created for each of the shared and private variants. For each breakpoint variant 3 probes of 120 bases each were created: Two reference probes leading up to the breakends from either side, as well as another SV probe going through the structural variant with the break junction close to the middle of the probe. The reference probes were designed to end 20 bases before the start of each structural variant breakend. The SV probe consists of any insert sequence flanked by equal number of bases from each side of the structural variant. For each single breakend variant 2 probes were created: One reference probe as described above and one SV probe which includes no more than 60 bases of the insert sequence with the remainder coming from the reference leading to the break point.

Together, this created a total of 17,125 capture probes of 120 nt in length, targeting 5,821 break-junctions (see supplementary table 1) which were ordered as custom target capture probes from Twist Biosciences (catalog ID 100533). For each of the 13 samples, 50 ng input DNA was used for indexed library construction with enzymatic fragmentation (Twist kit catalog IDs 100253, 100255 and 100401) according to the manufacturer’s protocol. A bead-based size selection was performed after PCR to remove the remaining larger fragments (>700bp).

Multiplexed hybridization was performed using the Twist Hybridization (ID100254), Blockers (PN100856) and Wash Kits (PN100214, 100215, 100216)) using Dynabeads™ MyOne™ Streptavidin T1 (Invitrogen PN65604D) following standard Twist protocol. Enriched library molecules were amplified by PCR for 11 cycles and sequenced on the Illumina NextSeq500 2x 150bp High Output run according to manufacturer’s standard protocol.

We created a set of predicted alternate contigs from the shared and private structural variant calls using the same technique from above for generating the (non-reference) SV probes and added these to the reference genome. We then mapped each of the reads from the capture panel output with BWA to a hybrid genome including both the GRCH37 reference genome and the novel alternate contig.

We assessed the viability of the probes by mapping each of the 120 base SV probes to the 2,000 base alternate contigs to determine its mapping quality. Of the 5,821 SV probes, 80 had 0 mapping quality and 5,377 had a perfect mapping quality of 60. Probes with a mapping quality of less than 20 were ignored as well as 77 micro-satellite probes that were unable to be unambiguously validated. Resultant BAM files were examined for evidence of the SV alternate contigs in the SV source sample BAM as well as the BAM files of each of the other samples as controls for systemic effects for each of the predicted variants. Specifically, the read depth on the alternate contig at the variant location was used to assess the validation status of the variant. SV calls were marked as validated if all the following criteria were met (and not validated otherwise):

- At least 2 reads were mapped to the alternate contig in the predicted sample
- The support rate for the alternate contig was significantly higher (Poisson model, p=0.001) in the predicted sample than the maximum of the other 13 samples
- <40 reads in total across all 13 control samples were mapped to the predicted alternate contig

### Comparison to PCAWG

Copy number data was obtained as for the Hartwig cohort running PURPLE ^2^ with default settings. PCAWG consensus SV and CN calls were obtained from https://dcc.icgc.org/releases/PCAWG/consensus_sv, and https://dcc.icgc.org/releases/PCAWG/consensus_cnv. Copy number transitions were matched with structural variants with a 100kb margin for PCAWG calls, and a 0bp margin for GRIDSS2/PURPLE. Copy number transitions in or within 100kb of centromeres or a gap in the reference genome were excluded from analysis. Copy number transitions matched by both a single breakend and a breakpoint, were considered breakpoint matches.

### Hartwig metastatic tumour cohort

GRIDSS2 was run on 3,782 paired tumour/normal samples from the Hartwig Medical Foundation cohort of metastatic solid cancers with a 32bp minimum event size. Samples were aligned with bwa against a GRCH37 reference genome containing only primary contigs. Single breakend RepeatMasker annotations were obtained by running *gridss*.*AnnotateInsertedSequence* against the UCSC GRCH37 (hg19) RepeatMasker track downloaded from http://hgdownload.cse.ucsc.edu/goldenpath/hg19/bigZips/hg19.fa.out.gz after converting to BED formart using bedops *rmsk2bed*.

Hartwig copy number was determined by PURPLE. Since PURPLE infers the copy number of short segments by the VAF of the flanking SVs, the copy number of these segments do not represent an independent validation of the SV. As such, FDR was segments from only the breakpoints in which the copy number of all 4 flanking segments was determined of depth of coverage and SNP BAF were considered.

Single breakends with a RepeatMasker annotation associated with centromeric or pericentromeric repeats were considered centromeric single breakends. The matching chromosome was considered to be the chromosome for which the BLAT^35^ based score score=(1000-((9-floor(Qsize/100))*mismatch+Qcount+Tcount))*min(match/Qsize,1)) is at least 900 and at least 25 higher than the best alignment on a different chromosome when aligning against hg38. A script for annotating likely centromere can be found in example/annotate_most_likely_centromere.R in the GRIDSS repository.

Phasability of the Hartwig cohort was calculated by determining, for each break junction, the length of the DNA segment if it was phased with the first break junction encountered in the appropriate orientation. Known phasing information was ignored for this analysis. Expected phasability was calculated by simulating 3,782 randomly fragmented paired genomes with the same number of break junctions as the corresponding Hartwig sample.

For both the PCAWG and Hartwig cohort, rearrangement event classifications were obtained by running LINX^28^ 1.12 on the GRIDSS/PURPLE outputs. Simulated FNR results were obtained by random subsampling of the Hartwig GRIDS2 SV calls and breaking LINX chains whenever a SV was excluded from the subsampling.

### COLO829 somatic benchmark

The COLO829T/COLO829BL cell lines (ATCC® CRL-1974™ and 1980™ respectively) were each sequenced three times to 100x/40x using the HMF workflow ^2,4^ and aligned against GRCH37 without alt contigs using BWA 0.7.15. GRIDSS 2.9.3, Manta 1.5.0, svaba 1.1.0, and smufin 0.9.3 ^36^ were run with default parameters. Programs were allocated 8,16,16,20 cores and 32, 32, 50, 500Gb of memory respectively. No smufin results were obtained in any replicate as smufin failed to complete in the 100,000 CPU hours/3 months wall time allocated.

Call matching was performed using the StructuralVariantAnnotation BioConductor package (DOI 10.18129/B9.bioc.StructuralVariantAnnotation). A 100bp matching margin was allowed around the break junction position. Tandem duplication calls matched with insertion calls if the size difference between the duplication and insertion was within 25bp. False positive calls under 50bp were filtered after matching so as not to penalise a caller reporting an event slightly larger than 50bp in the truth set, but slightly smaller than 50bp in the call set. If multiple calls in a call set matched a single truth set call, all except the highest QUAL call were ignored.

### COLO829 truth set generation

The COLO829 somatic SV truthset was generated using an orthogonal sequencing strategy. We sequenced the COLO829BL and COLO829T cell lines using Illumina HiSeqX (ENA run accessions ERR2752449 and ERR2752450 for COLO829BL and COLO829T, respectively), Oxford Nanopore (ERR2752451 and ERR2752452), Pacific Biosciences (ERR2752447 and ERR2752448) and 10X genomics (ERR2820166 and ERR2820167). All data is grouped under ENA study accession PRJEB27698.

Raw data was analysed for structural variants using the following tools:

- Illumina data was mapped using BWA 0.7.5, SV calling was performed with GRIDSS 2.0.1 and somatic SVs were filtered using gridss_somatic_filter.R.
- Nanopore data was mapped using NGMLR 0.2.6 and SV calling was performed with both NanoSV 1.2.0 and Sniffles 1.0.8 separately for COLO829T and COLO829BL. All SVs were
- merged with an overlap window of 200 base pairs using SURVIVOR 1.0.6 and SVs not present in COLO829BL were kept.
- Pacbio data was mapped using minimap2 2.11-r797 and SVs were called using pbsv 2.1.0 in joint calling mode for COLO829T and COLO829BL. Only SVs with no evidence in COLO829BL were kept.
- 10X genomics data was processed using Longranger 2.2.2 with default settings for COLO829BL and somatic mode for COLO829T. SV calls for both cell lines were merged with an overlap window of 200 base pairs using SURVIVOR 1.0.6 and SVs not present in COLO829BL were kept.

Somatic SV calls for each technology were merged with an overlap window of 200 base pairs using SURVIVOR 1.0.6. and all candidate breakpoints were subjected to independent validation by targeted capture and/or PCR-based approaches. SVs detected with two or more techniques that failed in these validation experiments were curated by manual inspection of the mapped reads using IGV ^37^. A total of 69 SVs were finally considered as true somatic SVs for COLO829T.

## Acknowledgements

S. E. de Garis for manuscript feedback and copyediting.

This publication and the underlying study have been made possible partly on the basis of the data that Hartwig Medical Foundation and the Center of Personalised Cancer Treatment (CPCT) have made available to the study.

## Author Contributions

DLC designed and implemented GRIDSS2. DLC, PP, JB, CS designed and performed dry lab experiments and analysis. JEV provided COLO829 golden reference data. NB performed independent break junction validation experiments. AH, RJ generated the GRIDSS2/PURPLE/LINX TCGA call set. ATP, PP, EC designed and supervised experiments. DLC, ATP, PP, EC contributed to writing of the manuscript.

## Availability of data and materials

GRIDSS2 source code is available as free and open source software from https://github.com/PapenfussLab/gridss under a GPLv3 license. GRIDSS2 releases are available as a github release, bioconda package, and docker image. Analysis scripts used to generate results are available from https://github.com/PapenfussLab/gridss/tree/master/scripts/gridss2_manuscript.

Hartwig cohort data was obtained from the Hartwig Medical Foundation (Data request DR-005). Standardized procedures and request forms for access to this data can be found at https://www.hartwigmedicalfoundation.nl/en.

Raw and analyzed data for the creation of the COLO829T/COLO829BL tumor/normal cell line pair structural variant truth set are available grouped under ENA study accession PRJEB27698. The COLO829 truth VCF is available from https://github.com/UMCUGenetics/COLO829_somaticSV.

Capture panel validations of 13 patient tumor samples are available under the controlled access dataset accession EGAD00001005525.

## Supplementary Figures

**Supplementary Figure 1:**
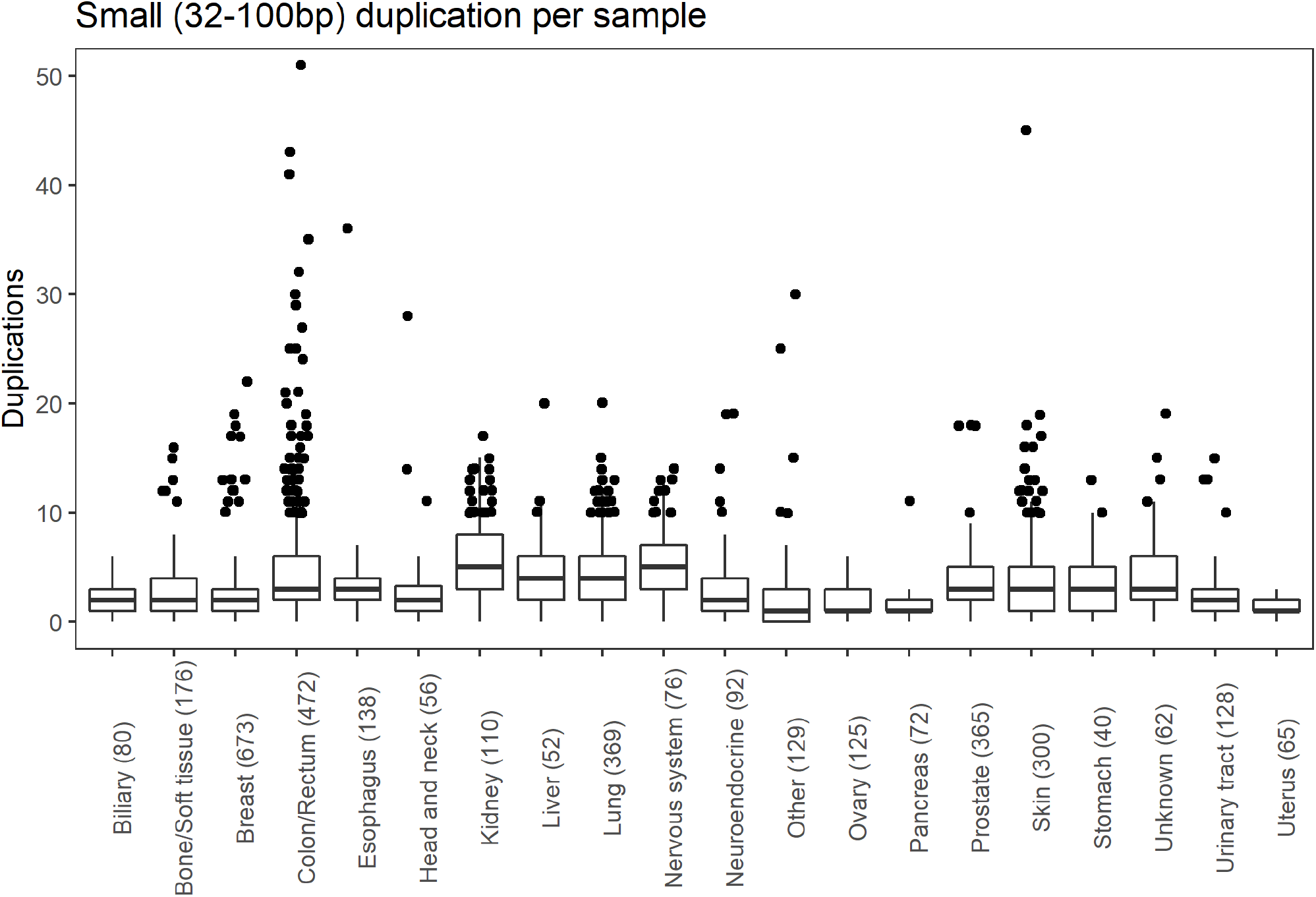
Distribution of 32-100bp duplication events per cancer type.

**Supplementary Figure 2:**
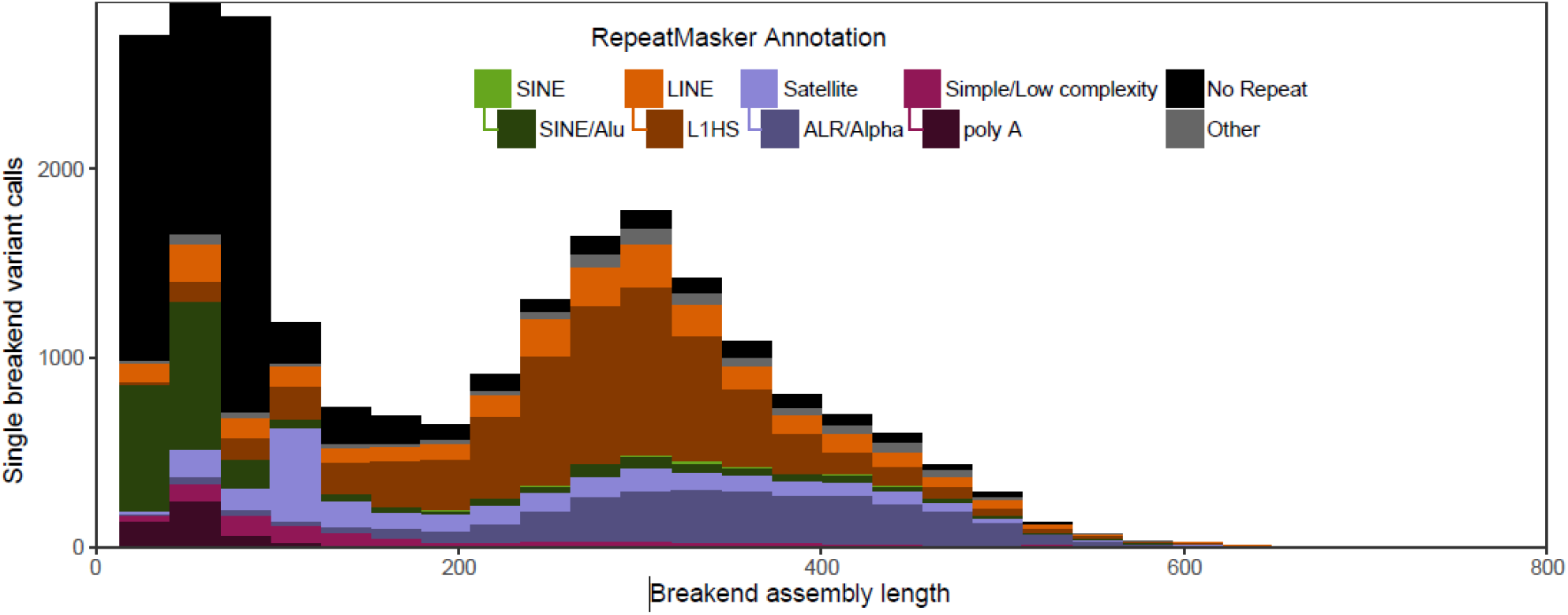
Single breakend RepeatMasker annotation for 1,528 PCAWG samples.

**Supplementary Figure 3:**
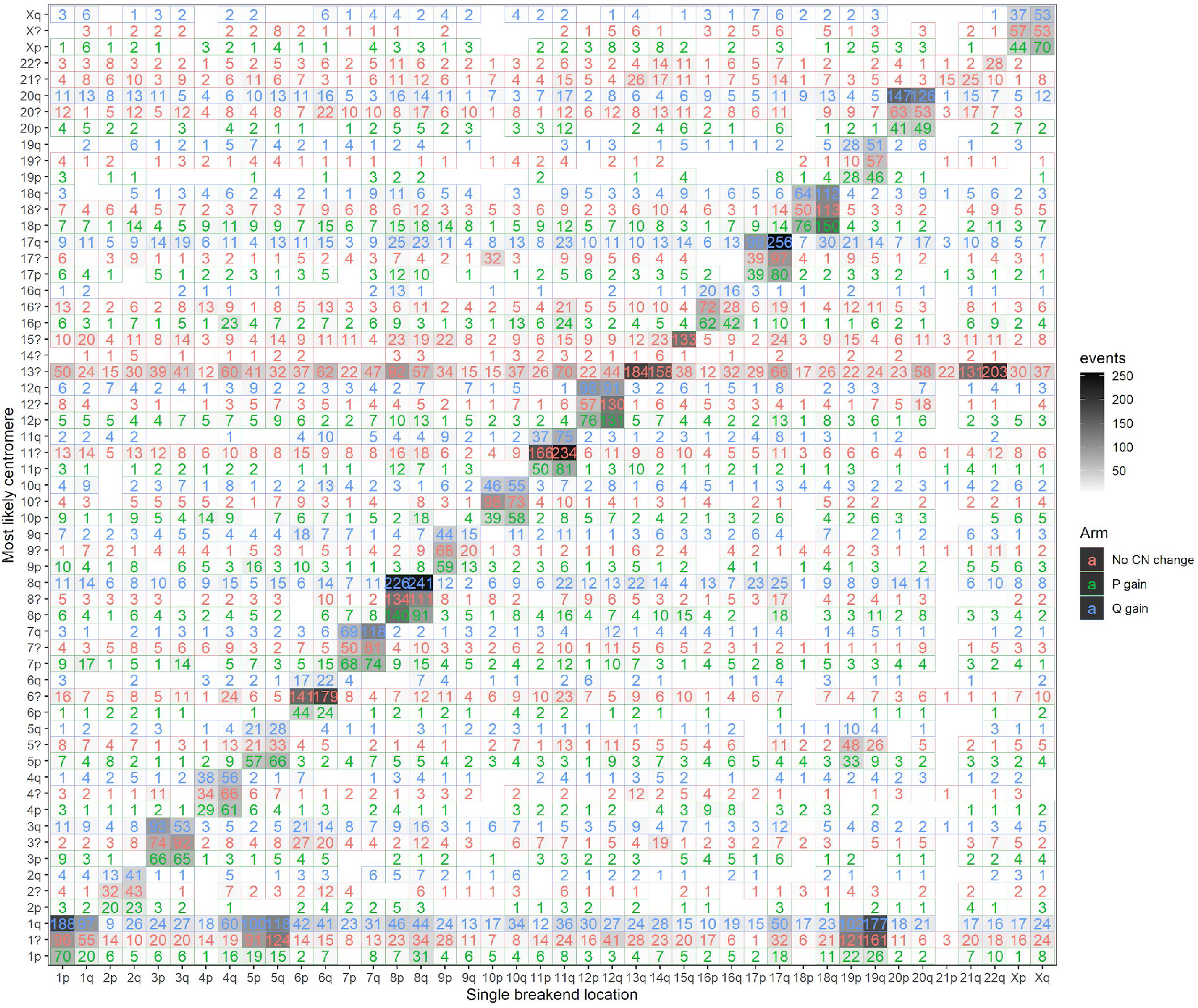
Heatmap of single breakends to centromeric sequence by chromosomal arm. The x axis indicates the arm of the single breakend with the y axis indicating the most likely chromosomal arm the single breakend is connected to based on the breakend sequence, and the copy number profile across the centromere. ? indicates the arm is unknown due to the lack of copy number change across the centromere, and p and q indicate a centromeric copy number gain to that arm.

**Supplementary Figure 4:**
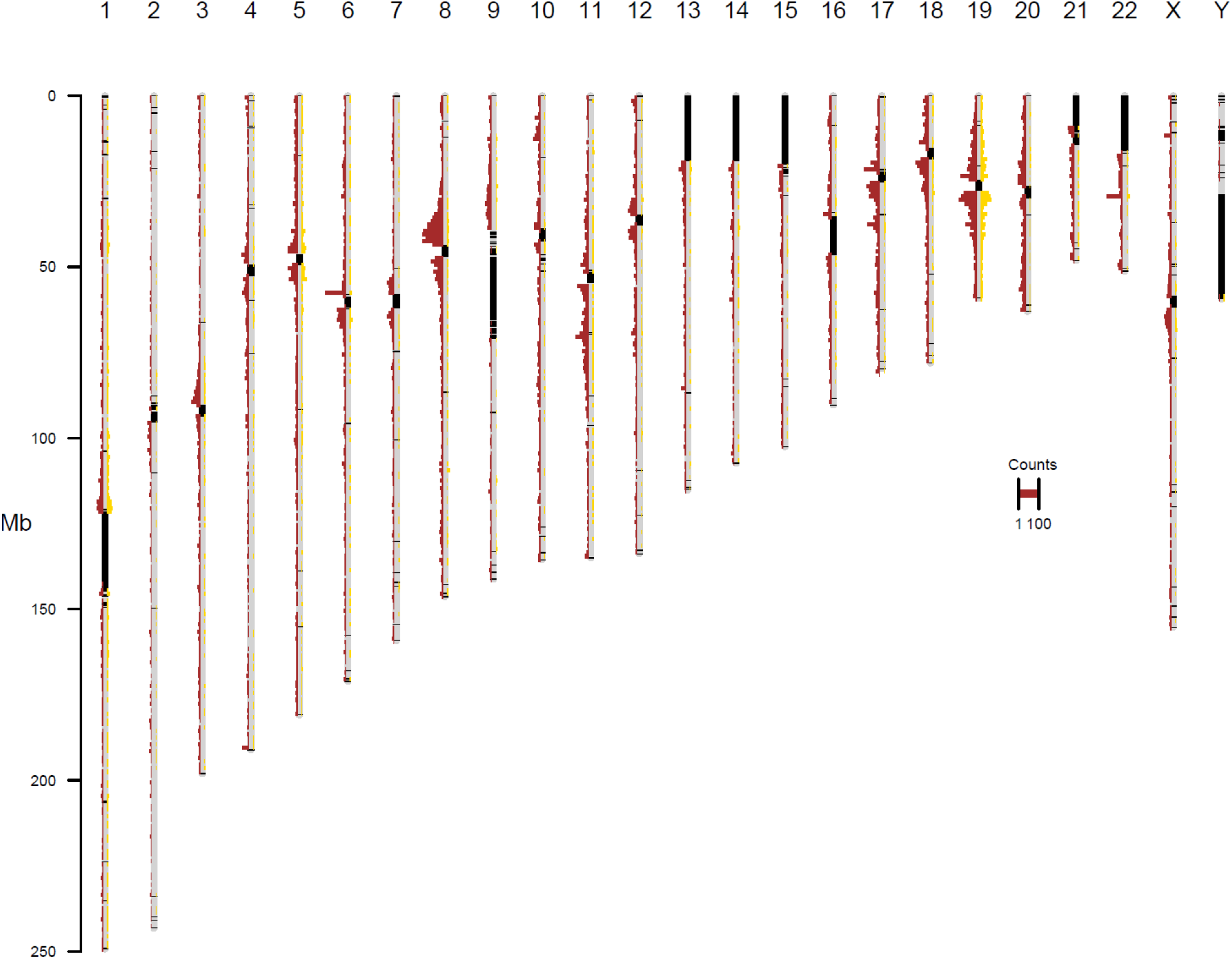
location of single breakends associated with the chromosome 1 centromere. Red indicates a single breakend associated with centromeric sequence on the same chromosome, yellow indicates a single breakend associated with centromeric sequence on chromosome 1. Single breakends to chromosome 1 occurring on chromosomes 5 and 19 follow a similar distribution to intra-chromosomal single breakends.

**Supplementary Figure 5:**
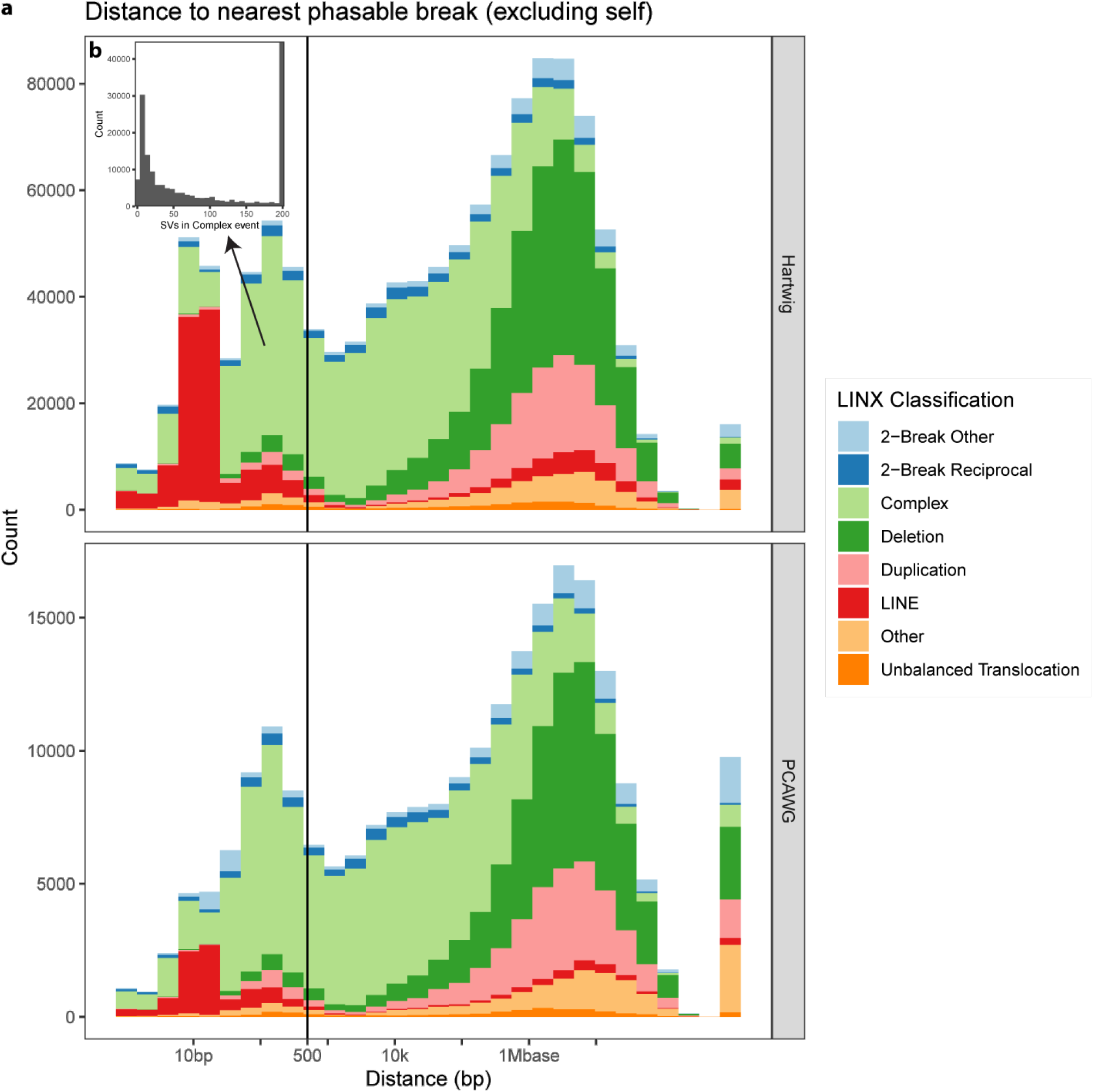
a) Phasability and LINX classification of structural variants in the Hartwig and PCAWG cohorts. Phasable variants are predominantly LINE translocation or form part of complex events. b) Number of SVs in complex event clusters containing phasable SVs. Most phasable SVs occur in highly complex events.

